# Suppression of Huntington’s Disease Somatic Instability by Transcriptional Repression and Direct CAG Repeat Binding

**DOI:** 10.1101/2024.11.04.619693

**Authors:** Ella W. Mathews, Sydney R. Coffey, Annette Gärtner, Jillian Belgrad, Robert M. Bragg, Daniel O’Reilly, Jeffrey P. Cantle, Cassandra McHugh, Ashley Summers, Joachim Fentz, Tom Schwagarus, Antje Cornelius, Ioannis Lingos, Zoe Burch, Marina Kovalenko, Marissa A Andrew, Frank C. Bennett, Holly B. Kordasiewicz, Deanna M. Marchionini, Hilary Wilkinson, Thomas F. Vogt, Ricardo M. Pinto, Anastasia Khvorova, David Howland, Vanessa C. Wheeler, Jeffrey B. Carroll

## Abstract

Huntington’s disease (HD) arises from a CAG expansion in the *huntingtin* (*HTT*) gene beyond a critical threshold. A major thrust of current HD therapeutic development is lowering levels of mutant *HTT* mRNA (m*HTT*) and protein (mHTT) with the aim of reducing the toxicity of these product(s). Human genetic data also support a key role for somatic instability (SI) in *HTT*’s CAG repeat – whereby it lengthens with age in specific somatic cell types – as a key driver of age of motor dysfunction onset. Thus, an attractive HD therapy would address both mHTT toxicity and SI, but to date the relationship between SI and HTT lowering remains unexplored. Here, we investigated multiple therapeutically-relevant HTT-lowering modalities to establish the relationship between HTT lowering and SI in HD knock-in mice. We find that repressing transcription of mutant *Htt* (m*Htt*) provides robust protection from SI, using diverse genetic and pharmacological approaches (antisense oligonucleotides, CRISPR-Cas9 genome editing, the *Lac* repressor, and virally delivered zinc finger transcriptional repressor proteins, ZFPs). However, we find that small interfering RNA (siRNA), a potent HTT-lowering treatment, lowers HTT levels without influencing SI and that SI is also normal in mice lacking 50% of total HTT levels, suggesting HTT levels, *per se*, do not modulate SI in *trans*. Remarkably, modified ZFPs that bind the m*Htt* locus, but lack a repressive domain, robustly protect from SI, despite not reducing HTT mRNA or protein levels. These results have important therapeutic implications in HD, as they suggest that DNA-targeted HTT-lowering treatments may have significant advantages compared to other HTT-lowering approaches, and that interaction of a DNA-binding protein and *HTT’*s CAG repeats may provide protection from SI while sparing HTT expression.

## Introduction

Huntington’s disease (HD) is a fatal neurodegenerative disease caused by the expansion of a glutamine-coding CAG repeat beyond a critical threshold in the *huntingtin* (*HTT*) gene ^1^. The clinical course of HD is characterized by progressive affective, cognitive, and motor disturbances, notably including chorea and other movement abnormalities ^2^. HD is inherited in an autosomal dominant manner, with disease fully penetrant beyond 39 CAG repeats ^3–5^. Onset of HD symptoms generally occurs in middle age, and age of motor dysfunction onset is highly influenced by the length of the CAG repeat, though significant variability in age of onset is observed across CAG lengths ^6^.

Somatic instability (SI), a further elongation of *HTT*’s expanded CAG tract in somatic tissues, has emerged as a key driver of HD pathogenesis. Evidence suggests that certain cell types, particularly striatal projection neurons, are particularly vulnerable to SI ^7–13^. The impact of SI in HD onset and progression is increasingly clear, thanks to large genome wide association studies (GWAS) that identified variants in DNA mismatch repair (MMR) pathway genes that bi-directionally alter age of motor onset in HD ^6,14,15^. Surprisingly, uninterrupted CAG repeat length also modifies age of onset - the “CAG tract” of most human *HTT* alleles contain a penultimate silent CAA interruption, and loss of this interruption leads to a markedly hastened onset of HD ^14,16^. Intriguingly, recent GWAS evidence confirms the robust impact of these onset-hastening loss of interruptions on the age of onset of HD, while contributing the novel observation that they are counterintuitively associated with significantly *reduced* SI, at least in blood ^15^. However, recent human single-cell RNAseq data supports a previously proposed model linking large CAG expansion to toxicity ^17^, claiming that toxic CAG lengths are much longer than inherited CAG lengths that cause HD, perhaps as large as 150 CAGs ^12^. Prior mouse genetic studies^18–20^ and other preclinical evidence also supports this model, as halting SI in HD knock-in mouse models with 140 CAG repeats provides dramatic protection from HD-relevant pathological signs ^21^, whereas the same intervention is ineffective in otherwise isogenic mice with ∼190-200 CAG repeats ^22^. An important question is which *HTT* gene product is the critical driver of toxicity at this CAG length. One proposal is an exon-1 only short *HTT* transcript variant (“*HTT1a*”) ^23^ and its encoded protein that avidly enters the nucleus, seeds aggregates ^24^, and is uniquely toxic compared to other mHTT fragments ^25^. Because the mis-splicing event that generates HTT1a is CAG size-dependent, a feasible model of toxicity is that SI drives CAG expansion in a given neuron to a length at which sufficient HTT1a is produced and the cells quickly sicken and die.

As a toxic gain-of-function disorder, there is broad interest in therapeutically reducing levels of the mutant *HTT* gene product(s) ^26^. The most clinically advanced effort is tominersen, an antisense oligonucleotide (ASO) developed by Ionis Pharmaceuticals and Roche that targets both *HTT* alleles and robustly reduced HTT levels in the cerebrospinal fluid of HD patients in a phase 1/2 study ^27^. Unfortunately, a subsequent phase 3 clinical trial of this ASO was halted early due to worsening trends in HD symptoms in ASO-treated patients; however, a new phase 2 study of the ASO to investigate different dosing strategies is underway (ClinicalTrials.gov ID NCT05686551) ^28^. Other therapeutic efforts to lower HTT include allele-selective ASOs targeting only m*HTT* by Wave Life Sciences, virally delivered micro-RNAs that reduce *HTT* by targeting exon-1 by uniQure (NCT04120493), and small molecule splice modulators that reduce levels of *HTT* transcript via selectively inducing nonsense mediated decay (NCT05358717) ^29^. Sangamo Therapeutics developed zinc finger transcriptional repressors (ZFPs) targeting expanded CAG repeats for allele selective silencing (now owned by Takeda Pharmaceuticals) ^30^. There are HTT lowering clinical and preclinical trials targeting both *HTT* alleles (Roche, uniQure, PTC, Skyhawk) and m*HTT* selectively (Wave, Takeda); that are capable of engaging HTT1a by targeting exon 1 (uniQure, Alnylam, Takeda) while others target downstream (PTC, Wave, Roche, Skyhawk). In terms of therapeutic modalities, these trials include viral gene therapy (uniQure, Takeda), oligonucleotides (Wave, Roche), and small molecules (PTC, Skyhawk).

We have a long-standing interest in HTT-lowering therapies ^31–33^. Motivated by a concern that HTT lowering may not be sufficient to address HD pathogenesis due to ongoing SI in *HTT*’s CAG tract ^34^, we set out to better understand the relationship between *HTT-*lowering treatments and SI *in vivo*. Specifically, we wanted to understand whether any *Htt-*lowering treatments in HD mouse models influence the progression of SI in *Htt*’s CAG tract. Using a range of genetic and pharmacological tools, we probed the links between *Htt-*lowering treatments, transcription, and SI. In general, we find DNA-targeted HTT-lowering therapies - notably ZFPs - consistently reduce SI, whereas those operating in the cytoplasm do not. Unexpectedly, we observe that a modified ZFP that can bind expanded CAG repeats, but not repress transcription, nevertheless robustly reduces instability, which suggests that delivery of CAG-binding DNA proteins may prove a novel means of developing therapies for HD by reducing SI without lowering HTT. An analagous DNA-binding protein approach could be envisioned for any repeat expansion disorder in which SI plays a role.

## Results

### Chronic ASO Treatment Reduces Somatic Instability and Slows Transcription in the Livers of *Htt^Q^*^111^*^/+^* Mice

In a previous study of the impact of peripheral silencing of *Htt* on HD central nervous system (CNS) phenotypes ^35^, we suppressed *Htt* in peripheral tissues of a knock-in HD mouse model with an inherited allele of ∼116 CAG repeats (*Htt^Q^*^111^*^/+^*) via weekly intraperitoneal administration of ASOs targeting exon 36 of mouse *Htt*. Because SI in the liver is as robust as the striatum, we wondered if this chronic HTT-lowering treatment might influence the progression of SI in this tissue ^36^. We found that treatment with a pan-*Htt*-targeting ASO (hereafter “*Htt* ASO”) from 2 to 10 months of age at 50 mg/kg/week reduced hepatic HTT levels by 64% at sacrifice ^35^; here, we observe that this is accompanied by a 37% reduction in the somatic expansion index of the *Htt^Q^*^111^ CAG tract in the liver (Fig. 1D-E). Somatic expansion in the striatum of these mice was unaffected by peripherally administered ASOs (Fig. S1, ANOVA treatment effect: F_(2,8)_ = 0.68, *p* = 0.53), which are excluded by the blood brain barrier ^37^. To confirm our observations, we generated two additional cohorts sacrificed after 5.5 or 12 months of treatment (Fig. 1A) – timepoints flanking our initial cohort’s treatment duration - in which we see comparable knockdown of HTT protein (Fig. 1B) and total *Htt* mRNA (Fig. 1C), and replicate our finding that decreased HTT levels are associated with decreased somatic expansion in the liver (Fig. 1D-E: N = 57 animals, 3-9 per treatment arm over three ages, ANOVA treatment effect F_(2,48)_ = 203.7, *p* < 2.2e-16). This reduction of expansion was not attributable to inherited allele sizes, which were similar across treatment groups (Table S2, ANOVA treatment effect F_(2, 54)_ = 0.42 *p* = 0.66). Increased duration of HTT suppression was associated with greater reductions in somatic expansion, with expansion indices reduced by 50% in the liver after 12 months of treatment (Fig.1E).

**Figure 1:**
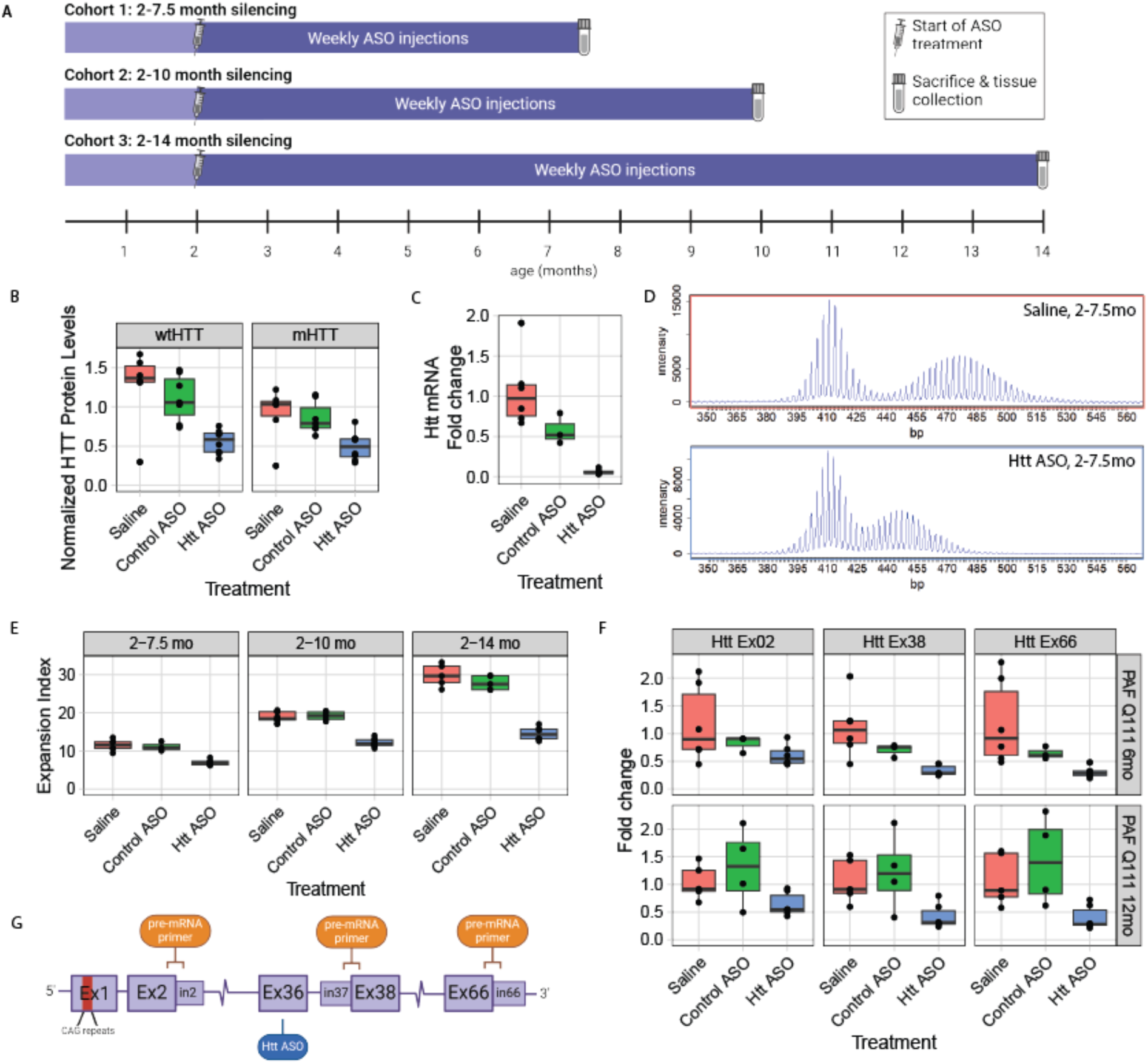
Chronic ASO treatment reduces somatic instability in mutant *Htt*’s CAG repeat and reduces *Htt* transcriptional rate in the livers of *Htt^Q^*^111^*^/+^* mice. **A)** Overview of peripheral ASO administration mouse cohorts. **B)** Chronic ASO treatment reduces levels of both wildtype (63% reduction; *p* = 0.001) and mutant HTT (54% reduction; *p* = 0.086) in the liver, as assayed with MSD at 14 months of age following 12 months of ASO treatment (N = 7-8 per arm). **C)** Total *Htt* mRNA, as assayed by qRT-PCR, is also reduced by chronic ASO treatment (94% reduction; p<0.0001). **D)** Exemplar traces of the size distribution of PCR products from a CAG-spanning PCR reaction. The top panel arises from a mouse treated with saline, while the bottom one is treated with *Htt*-targeted ASO. **E)** Somatic instability is reduced by chronic peripheral dosing of HTT ASO, regardless of length of treatment (39%, 35%, 51% reductions at 7.5, 10 and 14 months, respectively; *p* = 0.0002, *p* < 0.0001, *p* < 0.0001). **F)** Chronic ASO treatment reduces the levels of *Htt* pre-mRNA, as quantified by qRT-PCR with primer pairs that span the indicated exon-intron boundaries, with reduction increasing downstream of the ASO binding site (exon 2-intron 2: 44% reduction, *p* = 0.035; exon 37-intron 38: 66%, *p* = 0.009; exon 66-intron 66: 69%, *p* = 0.002). **G)** Overview of locations of the ASO target and exon/intron primer pairs. For box and whisker plots, each data point is shown, and the horizontal lines indicate the 25th, 50th and 75th percentiles of the data, with vertical lines indicating the range, with outliers detached from the vertical lines.

Recent publications have demonstrated that chronic ASO treatment can repress transcription in the nucleus via at least two distinct mechanisms ^38–40^, and that transcription through CAG repeats is known to influence SI *in vitro* ^41,42^. To quantify *Htt*’s transcription after ASO treatment, we established a qRT-PCR assay using exon-intron spanning primers designed to measure unspliced pre-mRNA at distinct points along the *Htt* transcript (Fig. 1F,G). Positive control experiments in cells treated with actinomycin D, a well-characterized transcriptional inhibitor ^43^, reveal dose- and time-dependent reductions in *Htt’*s pre-mRNA as detected with our pre-mRNA assay (Fig. S2). Applying this assay to the livers of mice chronically treated with ASO, where we observe significant reductions in SI, we find clear patterns of changes in the abundance of specific regions of the *Htt* transcript -- namely, robust reductions in pre-mRNA production at exons 38 (66% lower, *p* < 0.001) and 66 (69% lower, *p* = 0.003), downstream of the ASO binding site (Fig.1F). We note that even exon 2, 70kb upstream of the ASO binding site, had reduced production (44% lower, p = 0.035). Note that reported pairwise p-values refer to Tukey’s Honest Significant Difference (HSD) test conducted after factorial ANOVA, unless otherwise specified.

### Graded Transcriptional Repression with LacO Repressor Partially Rescues SI

To determine whether reducing transcription with a genetic system would also reduce SI, we quantified somatic expansion in mice in *Htt^LacO-Q^*^140^*^/+^*mice, in which expression levels of another knock-in CAG-expanded *Htt* allele (Q140) are regulatable via an upstream LacO repressor binding site (Fig. 2A) ^34,44^. This renders the *Htt^Q^*^140^ allele partially repressible by expression of a *LacI* repressor, driven by the broadly expressed β-actin promoter ^45^. Treatment with isopropyl β-d-1-thiogalactopyranoside (IPTG) recruits the lac repressor away from the targeted locus, enabling expression of the target gene. In our *Htt^Q^*^111^*^/+^* cohorts, we began ASO treatment at 2 months of age — before the emergence of most disease-associated phenotypes ^46^ and when low levels of somatic expansion are present ^47^. To investigate the relationship between duration of silencing and SI, we suppressed *mHtt* in *Htt^LacO-Q^*^140^*^/+^*mice starting at either embryonic day 5 or 2 months of age (Fig. 2B). We observe approximately 50% reduction in mHTT expression in the liver upon IPTG withdrawal (Fig. 2C), with consistent lowering of total *Htt* mRNA levels (Fig. 2D). Within each cohort, prolonging m*Htt* suppression led to greater reductions in somatic expansion of the targeted *Htt^Q^*^140^ allele (Fig. 2E, S3, ANOVA treatment effect at 6 months of age F_(2,86)_ = 148.3, *p* < 2.2e-16; at 12 months of age F_(2,123)_ = 91.1, *p* < 2.2e-16; exemplar PCR product traces in 2F). The reduced expansion indices did not reflect differences in inherited allele sizes, which were similar across groups (Table S2). Consistent with our ASO data, we find that allele-selective repression of the *mHtt* allele is effective to significantly lower SI.

**Figure 2:**
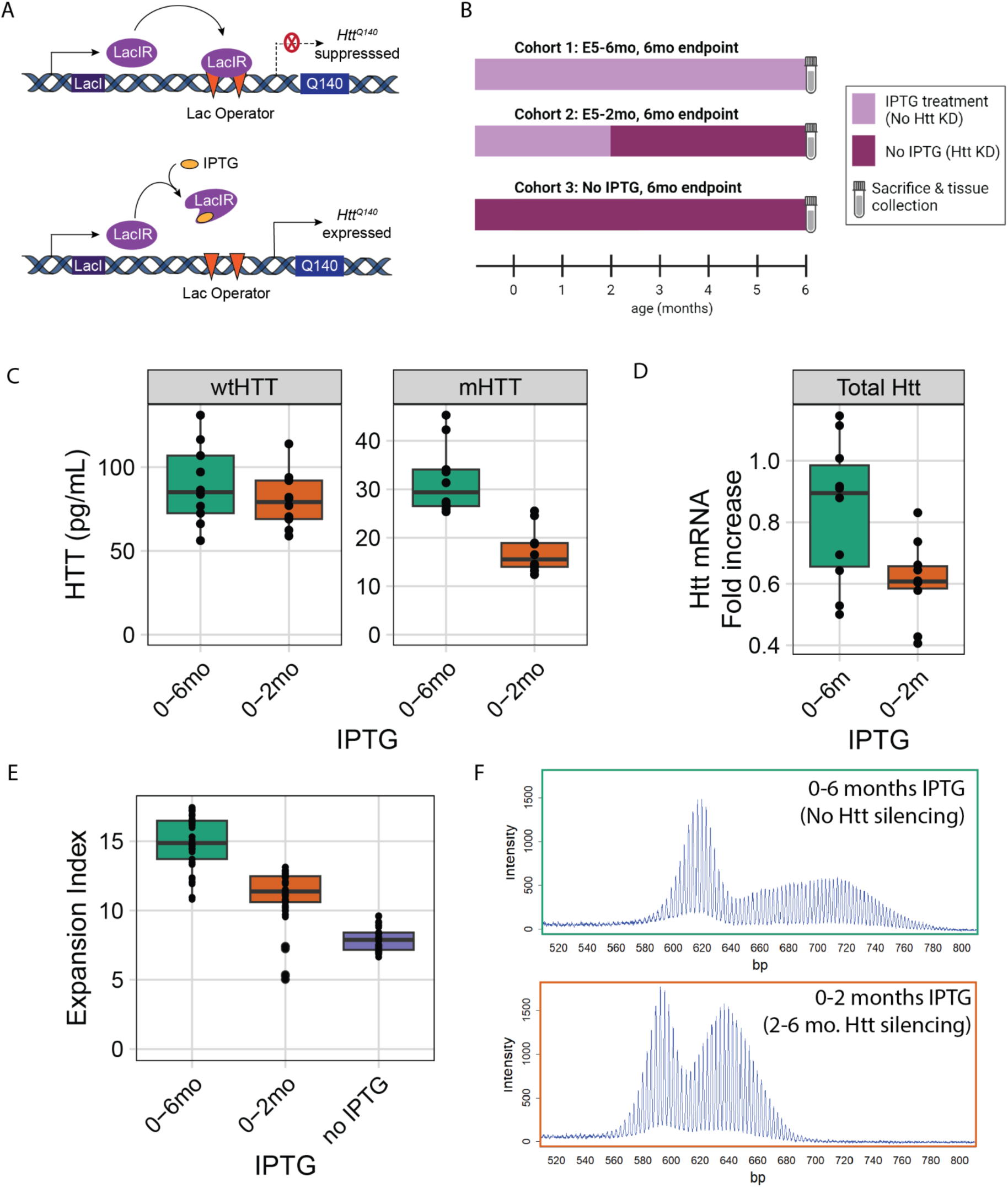
Partial *lac* repressor-mediated transcriptional repression of m*Htt* reduces somatic instability in *Htt*’s CAG repeat in the livers of *Htt^Q^*^140^ mice. **A)** Schematic of study design indicating when mice in 3 cohorts were treated with IPTG, reducing LacR-mediated repression of the *Htt^Q^*^140^ allele. **B)** Overview of the mechanisms of action of the *lac* repressor system. **C)** Protein knockdown at sacrifice: wtHTT is unaffected by the lac repressor (*p* = 0.52), while mHTT levels are reduced (46%, *p* < 0.0001). **D)** Knockdown of total *Htt* mRNA from liver from replicate mouse experiment (27%, *p* = 0.018; NB, only mHTT is repressed, so this non-allele selective mRNA assay represents an under-estimate of mHTT-selective knockdown). **E)** Somatic instability is reduced via partial LacR-mediated repression of m*Htt*, with greater suppression of instability given longer suppression durations. **F)** Exemplar traces of repressed and unrepressed mice. For box and whisker plots, each data point is shown, and the horizontal lines indicate the 25th, 50th and 75th percentiles of the data, with vertical lines indicating the range, with outliers detached from the vertical lines.

### CRISPR/Cas9-mediated Excision of Proximal Promoter Regions Protects from SI

To establish the impact of more complete loss of *mHtt* transcription on SI in adult animals, we devised a genome editing approach to excise the proximal promoter region and start codon in *Htt^Q^*^111^*^/+^* mice. We reasoned that by selectively removing the promoter regions ^48^ and start codon from the *Htt^Q^*^111^ allele, while sparing the CAG repeats, we could establish the relationship between adult-onset, complete m*Htt* silencing and SI. We designed and validated paired guide RNAs (gRNAs) targeting unique regions of sequence in the *Htt^Q^*^111^ allele - one in the 5’ untranslated region upstream from *mHtt*’s start codon, and a second in exon-1 of *mHtt*, just upstream of the CAG repeat (Fig. 3A). We reasoned that successful dual cutting by Cas9 and joining the double strand break ends by non-homologous end joining would eliminate the required transcriptional regulatory elements driving *mHtt* expression, whilst sparing the CAG repeats ^49,50^. We developed a single plasmid to express a fluorescent protein and two gRNAs, flanked by adeno-associated virus (AAV) inverted terminal repeats to enable AAV generation. Using this plasmid, we generated AAV8, which has very high tropism for hepatocytes ^51^, screened 8 gRNA pairs in mice, and chose a pair that resulted in potent, allele-selective HTT lowering (Fig. 3B). This gRNA pair is referred to as “QPro” (Q111 promoter) in the figures below. *Htt^Q^*^111^*^/+^; Cas9* mice were treated via tail vein injection at 2.5 months of age and sacrificed at 8 months of age. AAV8::QPro expression resulted in robust reduction of mHTT protein levels, while sparing wtHTT (Fig. 3B). Supporting our previous results in transcriptional repression of *mHtt*, preventing transcription via QPro treatment resulted in robust reductions in SI compared with expression of the empty vector (Fig. 3C-D). We recognize that the robust impact of editing on SI may arise from multiple cis-mediated effects, such as altered chromatin conformation, with the common downstream impact of reducing SI.

**Figure 3:**
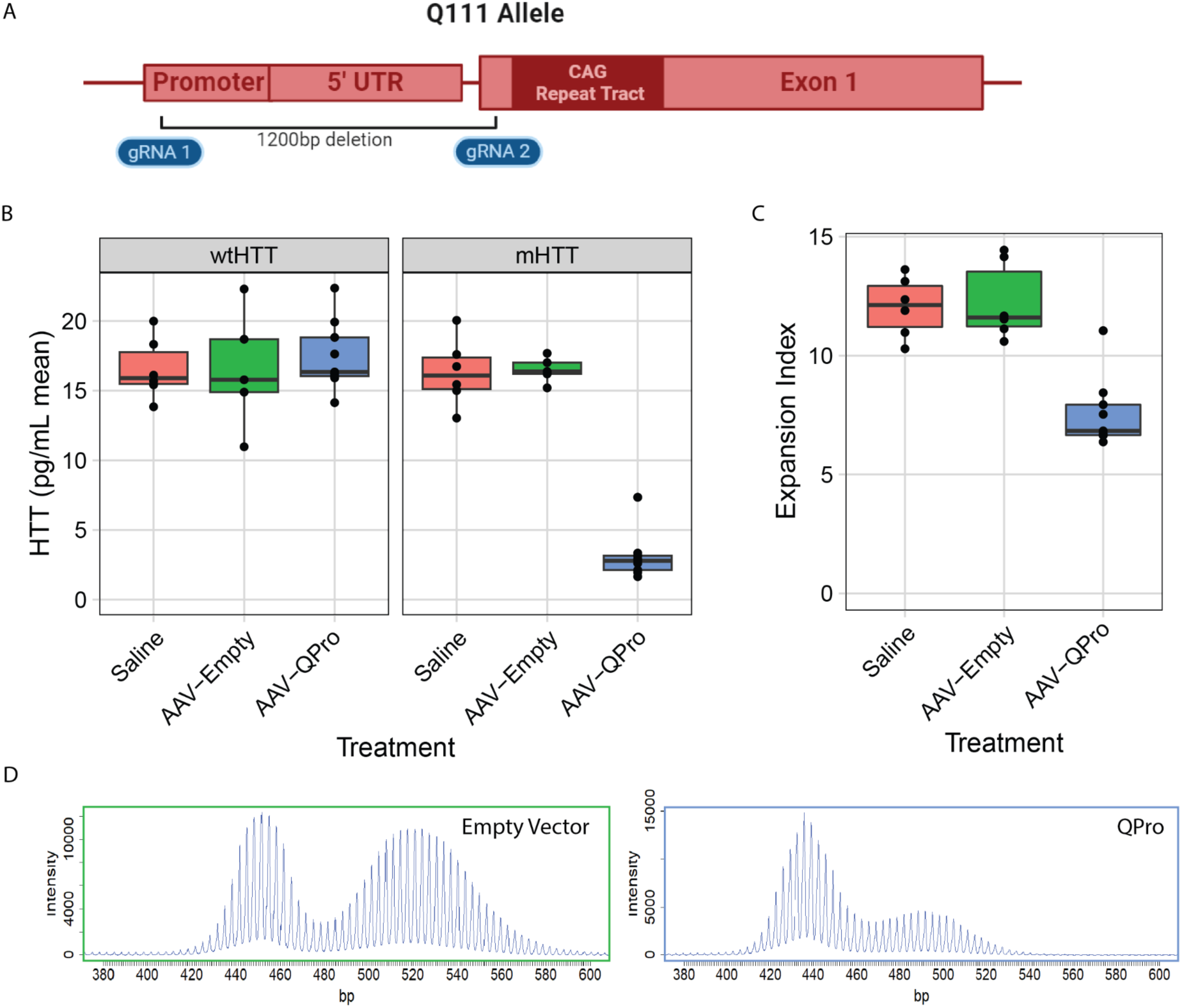
CRISPR/Cas9-mediated deletion of the m*Htt* promoter region reduces SI in *Htt*’s CAG repeat tract. **A)** Cartoon indicating the location of the gRNA pair referred to as “QPro,” spanning an approximately 1,200bp region, that targets unique sequences in the humanized *Htt^Q^*^111^ allele that are not present in mouse *Htt*. **B)** AAV8-delivered treatment with QPro results in robust, allele-selective lowering of HTT protein in the livers of *Htt^Q111+^*; Cas9 ^73^ mice. **C)** Treatment with AAV8::QPro at 2.5 months of age results in reduction of expansion index at 8 months of age. **D)** Exemplar traces of CAG tract expansions in untreated and treated animals.

### Chronic Lowering of HTT with siRNA Does Not Influence SI in the liver

The previous experiments provide strong support that suppressing *Htt* transcription reduces somatic expansion. However, the results do not rule out the possibility that HTT protein could act *in trans* to promote repeat expansion, with the reduction of somatic expansion occurring indirectly as a result of HTT protein loss. To test this, we first determined whether nuclear access is required for a HTT-lowering treatment to affect SI by using small interfering RNA (siRNA), a silencing method that acts only in the cytoplasm, to target *Htt*. Since siRNA is enacted by the cytoplasmic RISC complex ^52^, we reasoned that lowering *Htt* via siRNA would be unlikely to directly influence transcription in the nucleus. We treated a cohort of *Htt^Q^*^111^*^/+^* mice from 2.5 to 6 months of age with monthly injections (10 mg/kg) of a well-characterized pan *Htt*-targeting siRNA (HTT10150) ^53^ conjugated to N-acetylgalactosamine (GalNAc), which facilitates specific uptake in hepatocytes (Fig. 4A) ^52^. This treatment markedly reduced both wildtype and mutant HTT protein (Fig. 4B, wtHTT 74% lowered, mHTT 55% lowered) and total *Htt* mRNA levels (Fig. 4C) in the liver. However, siRNA treatment did not impact transcription as determined by our pre-mRNA assay (Fig. 4D). Consistent with our broader hypothesis, this chronic, robust lowering of HTT in the liver also did not alter SI in *Htt*’s CAG tract (Fig. 4E), confirming that HTT levels *in trans* do not regulate somatic instability.

**Figure 4:**
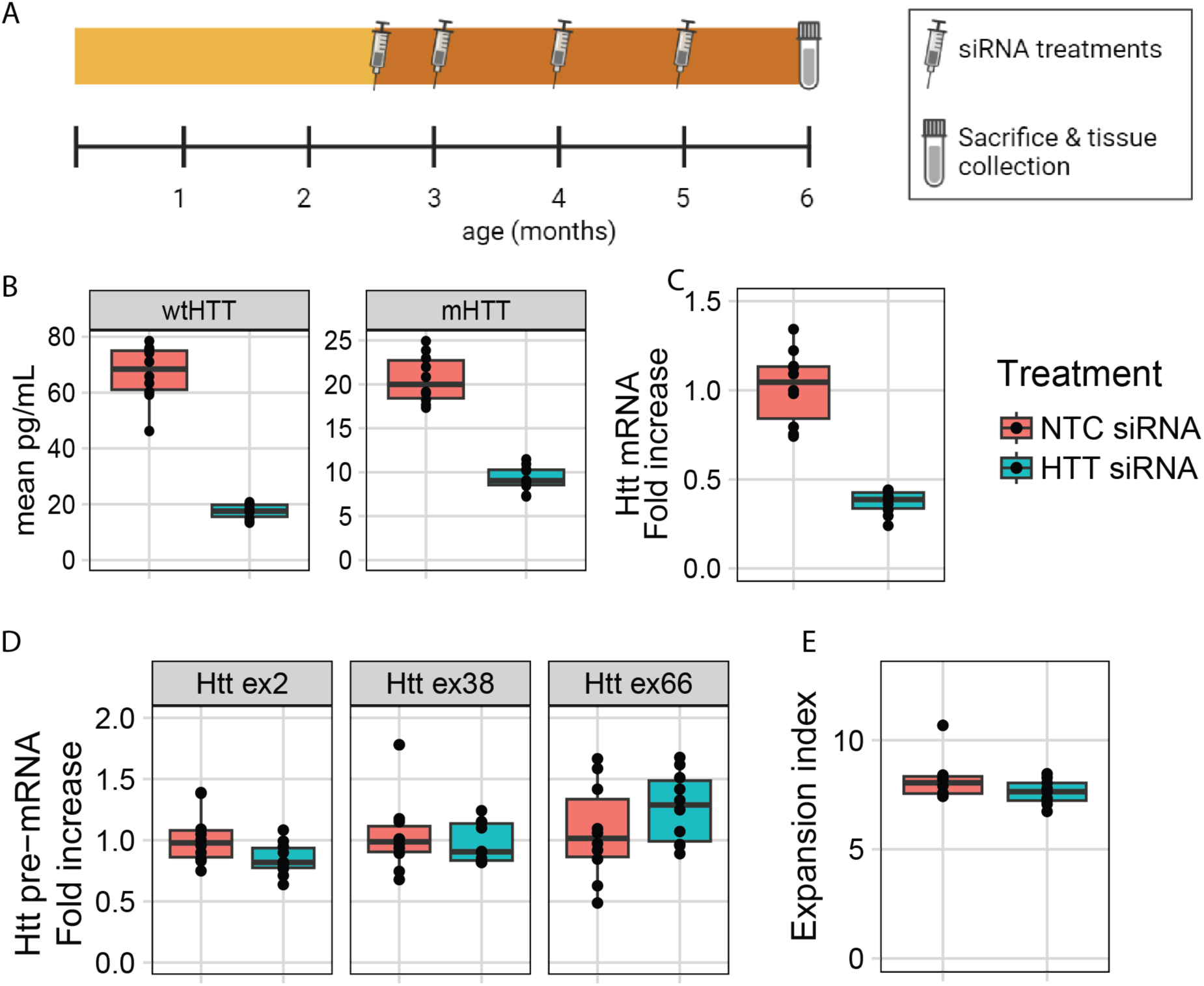
siRNA-mediated repression of *Htt* does not alter its somatic instability or transcriptional rate in *Htt^Q^*^111^*^/+^*mice. **A)** Schematic of study design. **B)** Chronic (10 mg/kg/month) siRNA treatment reduces levels of both wildtype (74% reduction; *p* < 0.0001) and mutant HTT (55% reduction; *p* < 0.0001) in the liver, as assayed with MSD at 6 months of age following 3.5 months of treatment (N = 10 per arm). **C)** Total *Htt* mRNA, as assayed by mouse *Htt* qRT-PCR, is also reduced by siRNA treatment (64% reduction; N = 10, *p* < 0.0001). **D)** *Htt* pre-mRNA levels are not impacted by siRNA treatment (NS). **E)** Chronic siRNA treatment does not change SI in HTT’s CAG tract (NS).

### Elimination of Wildtype HTT has no Impact on SI in the Brain and Liver

As an additional test of whether wildtype HTT might act *in trans* to influence SI, we generated *Htt^Q^*^111^*^/-^* hemizyous knock-in mice in crosses between *Htt^Q^*^111^ and *Htt*^ex4/5^ knockout mice, expressing one CAG-expanded allele and one *Htt* null allele (*Htt^Q^*^111^*^/-^*) ^54^. Western blotting confirmed the loss of wildtype HTT and consistent levels of mHTT expression in *Htt^Q^*^111^*^/-^* compared to *Htt^Q^*^111^*^/+^* mice (Fig. 5A-B). We found no differences in somatic expansion in the liver or striatum between *Htt^Q^*^111^*^/+^* and *Htt^Q^*^111^*^/-^*mice, matched for inherited repeat length, at 3, 6 and 10 months of age (Fig. 5C). Thus, 50% reduction of HTT levels (and all wildtype HTT) has no impact on SI in m*Htt*’s CAG repeats, providing further evidence that HTT protein does not act *in trans* to modify SI. This lack of effect is consistent with previous phenotypic characterization of HD-relevant phenotypes in *Htt^Q^*^111^*^/-^* mice, demonstrating that loss of wildtype HTT does not impact nuclear accumulation of mHTT ^55^.

**Figure 5:**
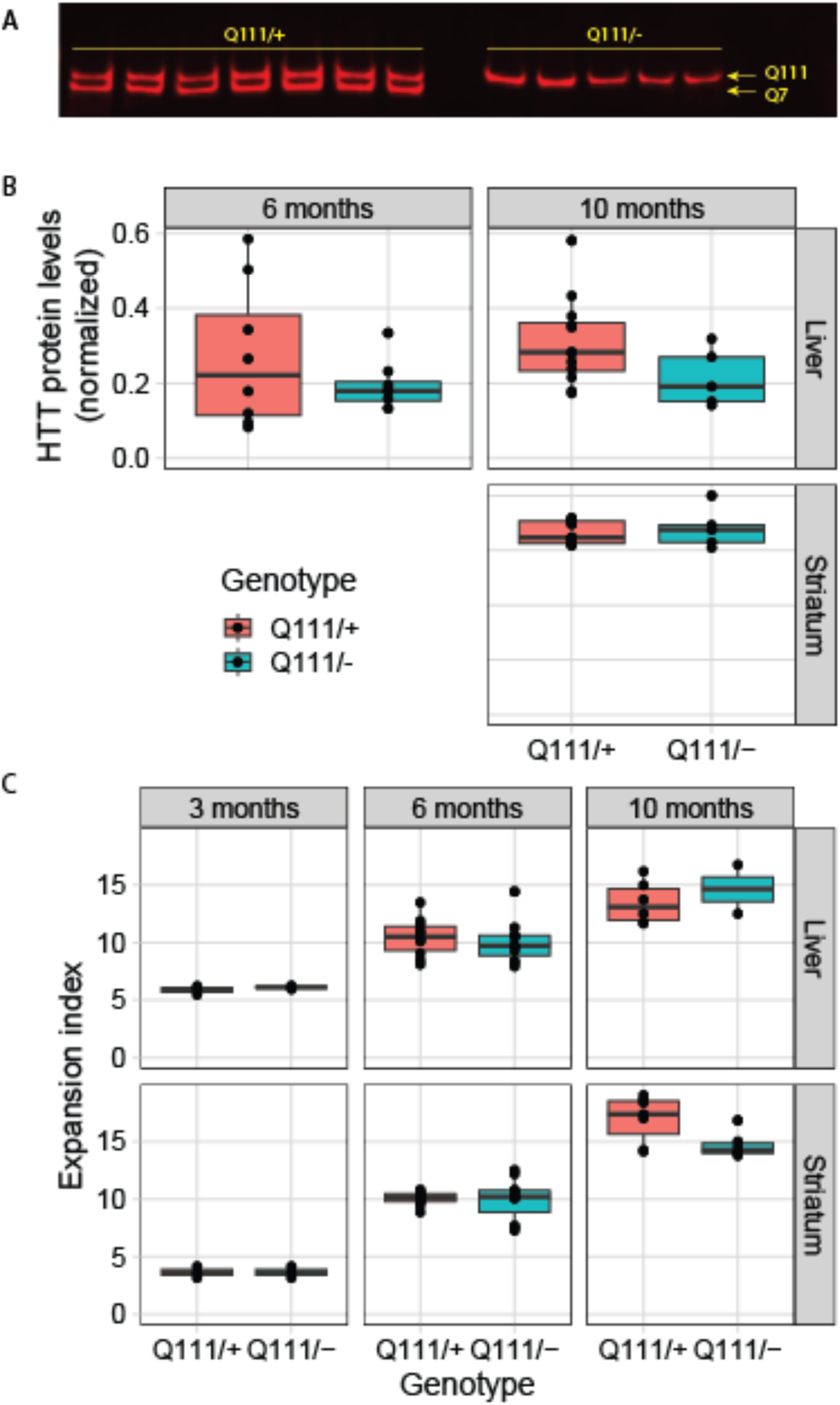
Genetic loss of wtHTT does not influence somatic instability. **A-B)** Western blot demonstrating the lack of wildtype HTT in *Htt^Q^*^111^*^/-^*mice, and no alteration in mHTT levels. **C)** Instability in *Htt*’s CAG repeat is not altered in *Htt^Q^*^111^*^/-^* mice compared to *Htt^Q^*^111^*^/+^* mice in either the striatum or liver.

### ZFP-mediated transcriptional repression and CAG repeat binding both reduce SI

Taken together, our data point to SI being directly influenced by *Htt* transcription itself. To further interrogate this hypothesis using an orthogonal therapeutic paradigm, we turned to well-characterized ZFPs targeting *mHtt* ^30^. These selectively bind *Htt*’s expanded CAG repeat via a zinc finger DNA binding domain (DBD) fused to a Krüppel-associated box (KRAB) domain that induces repressive chromatin around *Htt*’s start codon. To directly examine the relationship between CAG binding, transcription and SI, we engineered ZFPs lacking the DNA binding domain (referred to as KRAB-only) or the KRAB domain (referred to as DBD-only) (Fig. 6A). We packaged the three resulting constructs into AAV1/2 serotype capsids and injected them directly into the striatum of *Htt^Q^*^111^*^/+^*mice. Using confocal microscopy, we confirmed widespread, consistent delivery of all three constructs to injected striata. Staining for an epitope tag delivered by the construct shows that full ZFP-KRAB, but not DBD-only or KRAB-only, is anti-correlated with the formation of nuclear aggregates of mHTT, as has previously been reported ^30^ (Fig.S4A). This confirms efficient delivery of our ZFP constructs to injected striata, and that full ZFP treatment robustly reduces aggregated *mHtt* species in the nucleus.

**Figure 6:**
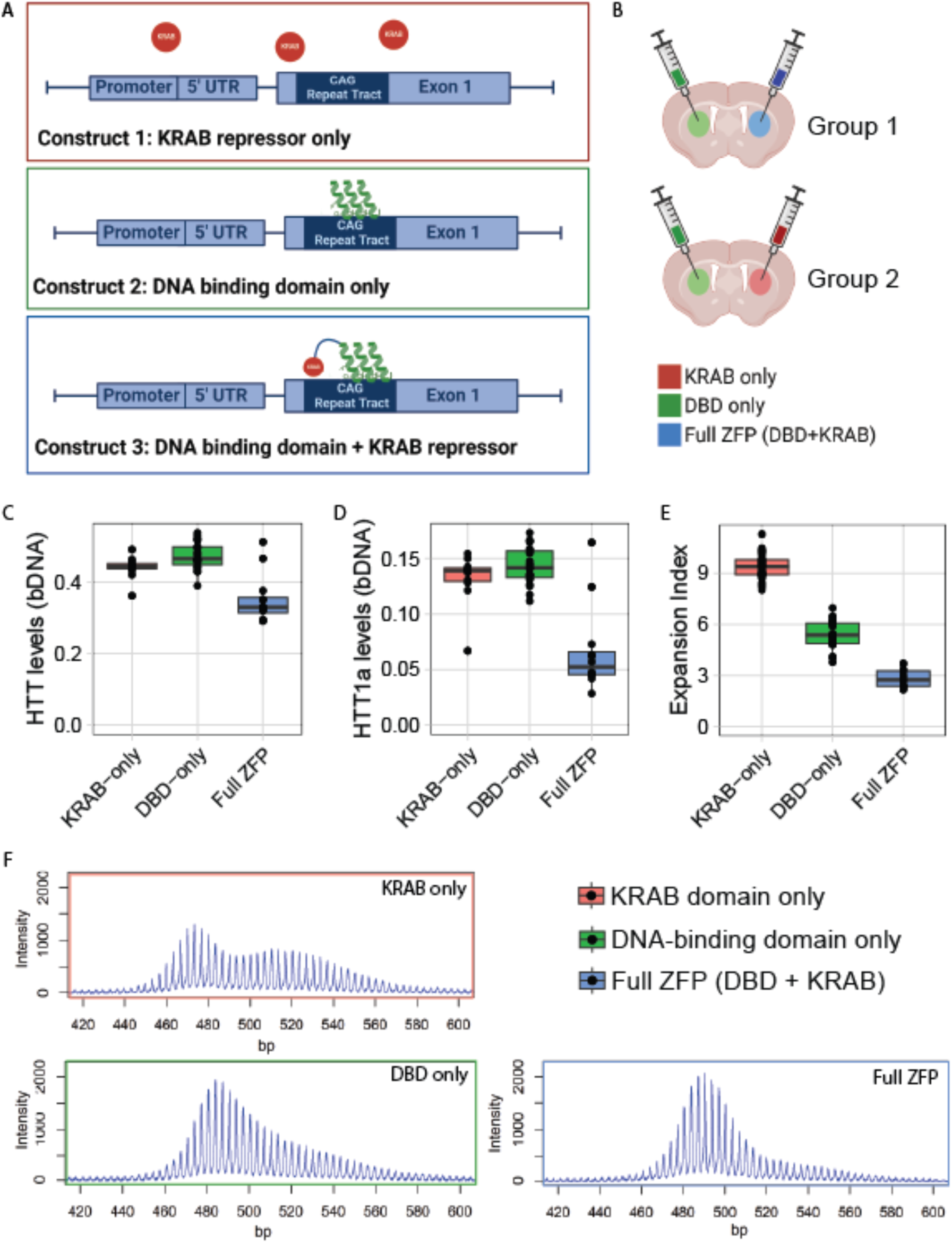
ZFP treatment reduces HTT’s transcriptional rate and blocks CAG SI, while delta-KRAB lowers SI but spares transcription. **A)** Schematic of ZFP constructs: the DNA-binding domain (DBD) binds CAG repeats, while the KRAB domain induces repressive chromatin. **B)** Schematic of treatment paradigm - each subject mouse is treated with two viruses - one in each hemisphere. **C)** *Htt* transcript levels as measured by bDNA assay decrease after ZFP treatment (full ZFP vs KRAB-only, 20% reduction, *p* = 0.0002) but are unaffected by DBD-only treatment (NS). NB, this is a non-allele selective assay, and so likely underestimates the amount of allele-selective lowering of m*Htt*. **D)** Treatment with the complete ZFP constructs leads to a marked reduction in exon-1 only m*Htt* (*Htt1a*) levels as measured by bDNA assay (50% reduction, *p* < 0.0001), but are unaffected by DBD-only treatment (NS). **E)** Somatic instability of m*Htt*’s CAG tracts is reduced (70%, *p* < 0.0001) by the full ZFP construct, with the DBD-only construct resulting in an intermediate reduction (42%, *p* < 0.0001). **F)** Exemplar traces from each treatment group.

**Figure 7.**
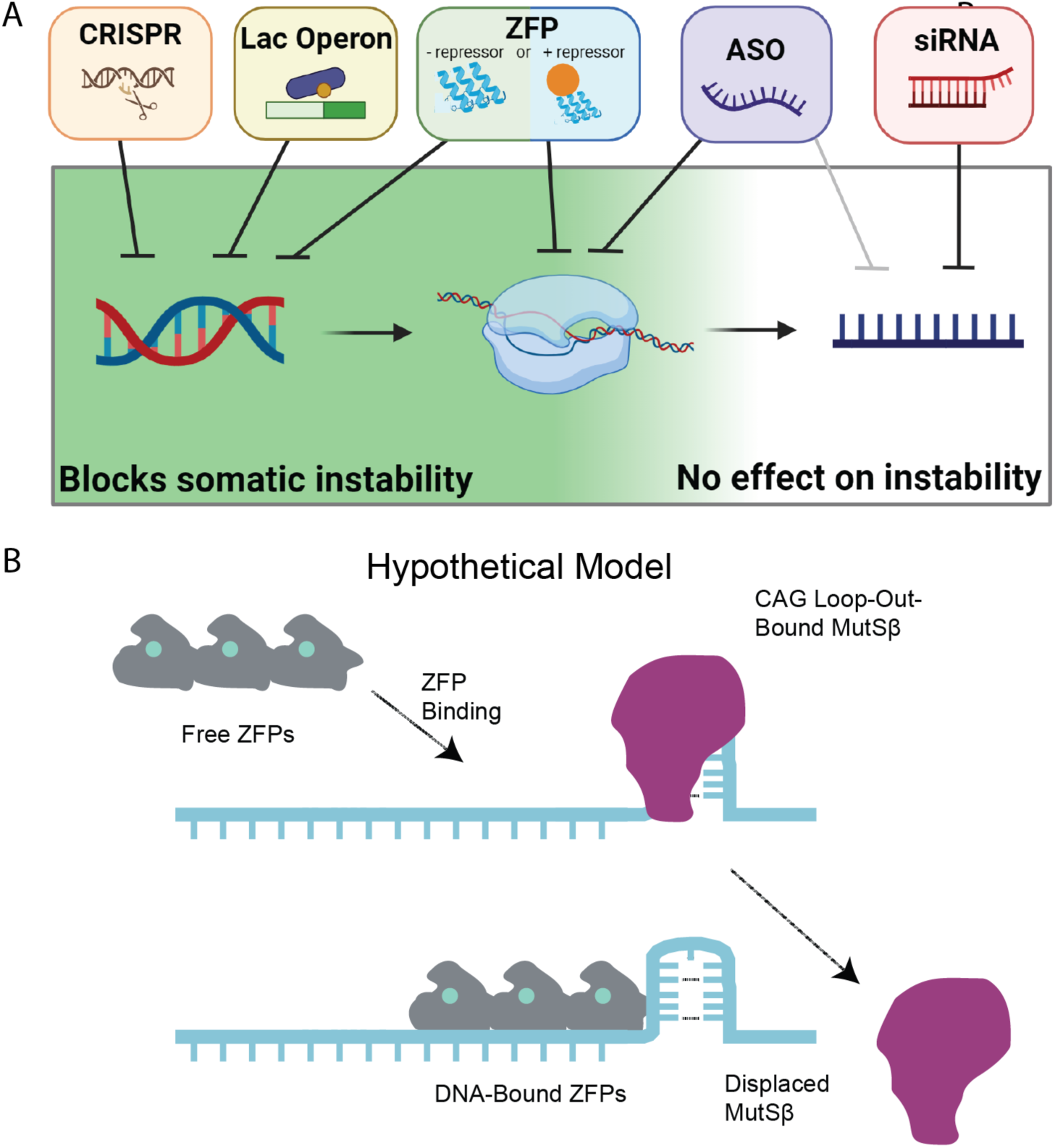
graphical abstract: Summary of our findings of the impact of HTT-lowering treatments on somatic instability.

We next investigated somatic instability and *Htt* transcript levels in *Htt^Q^*^111^*^/+^*mice after intra-striatal ZFP injections at 2 months of age. Each mouse was injected with one viral construct in one hemisphere, and another construct in the contralateral hemisphere, allowing us to make within-mouse comparisons of the impact of ZFP treatments (Fig. 6B). At 6 months of age, we collected striatal tissue and quantified *Htt* transcripts and SI in *Htt*’s CAG tract.

Compared to the DBD-only control, treatment with the full ZFP construct reduced total *Htt* transcript levels (20% reduction; Fig. 6C, *p* = 0.0002), consistent with previous observations with these reagents ^30^, and our preliminary studies (Fig. S4A-D). The DBD-only version has no impact on full-length *Htt* levels (Fig. 6C, *p* = 0.14), confirming that the KRAB domain is required to repress transcription. We also examined levels of an exon 1-only isoform of *Htt*, referred to as *Htt1a*, arising from aberrant splicing of exon-1 when CAG repeats are expanded (Fig. 6D) ^23^. In wildtype mice, no appreciable levels of this isoform are generated at 6 months of age ^56^, so the ZFP-sensitive *Htt1a* levels seem likely to arise from the m*Htt* allele. *Htt1a* levels decrease by 50% after treatment with full ZFP construct (*p* < 0.0001), and are unaffected by DBD-only (*p* = 0.44). This is in accordance with our observed 20% *Htt* reduction; since that assay is not allele-selective, it should detect about half the level of knockdown of the *Htt1a* assay. We take this as supportive evidence that m*Htt* transcription was reduced by the full ZFP, but not the DBD-only construct. Additionally, we investigated the effects of the different ZFP constructs on a number of predicted off-target genes. The full ZFP lowers off-target transcripts of *Atp2b3*, *Dnajc12*, *Eid2b*, *Nap1l3*, and *Stk32c*; however, the DBD-only construct does not affect levels of any predicted off-target transcript (Fig. S4C). This provides further evidence that the DBD-only construct cannot suppress transcription.

Finally, we examined the impact of all 3 ZFP treatments on SI. Strikingly, treatment with the DBD-only ZFP has a marked impact on SI (42% reduction, Fig. 6E-F, *p* < 0.0001), despite not repressing m*Htt* transcription. Even greater reductions in SI were achieved after delivery of the full transcriptionally repressive competent ZFP+KRAB (70% Reduction, Fig. 6E-F, *p* < 0.0001). This likely under-estimates the amount of protection provided, given that transduction efficiency is not 100%, and at treatment initiation at 2 months of age some SI has occurred. To confirm that the DBD-only construct does not affect transcription, we measured nascent mRNA in STHdhQ111 cells treated with each of the three ZFP constructs. We then amplified pre-mRNA from *Htt* exons 36 and 66. We find that pre-mRNA is reduced only by the full ZFP, and importantly, is unchanged by the DBD-only construct (Fig. S4D). This provides further evidence that binding CAGs alone, without any effect on *Htt* transcription, can provide protection from SI.

## Discussion

Our study demonstrates that repressing transcription or binding to *HTT’s* CAG tract in HD mouse models significantly reduces SI *in vivo*. Potently lowering m*Htt* with siRNA, which acts in the cytoplasm modulates neither transcription nor SI. We also demonstrate that CAG-targeting ZFPs reduce HTT1a levels, making them, to our knowledge, the only HD therapeutic to: allele-selectively lower mHTT, reduce HTT1a levels, and block instability. As such, we believe that epigenetic editing techniques such as ZFPs have great theoretical advantages as therapies for HD. Intriguingly, we find that the DBD-only version of our ZFP significantly reduces SI, despite not lowering HTT levels. We speculate that this may be by a steric block preventing access to CAG loop-outs that, when bound by the MutSβ complex, initiates the chain of events leading to SI ^57^. Another possible explanation is that the DBD itself stabilizes CAGs, reducing the occurrence of loop-outs. Testing of this model is ongoing, and if accurate, suggests that non-transcriptionally-repressive DNA-binding proteins are a novel means to block SI without lowering the levels of the targeted protein. This may be particularly important in disorders for which knockdown of the host protein would be poorly tolerated, such as SCA17, which is caused by a CAG repeat expansion in the TATA binding protein (TBP) ^58^. Indeed, the CAG repeat-targeted ZFPs we tested here could immediately be applied to the nine known CAG repeat disorders (SBMA, DRPLA, HD, SCA -1, -2, -3, -6, -7 and -17) ^59^.

Active transcription increases SI of various genomic repeats in diverse model systems (reviewed in ^60,61^). Compelling data have emerged from the study of fragile X-related disorders, which arise from CGG expansions in the X-linked *FMR1* gene ^62^. The Usdin lab has demonstrated that X-chromosome inactivation stabilizes *FMR1*’s CGG repeat in mouse models, consistent with transcription being an important factor controlling instability *in vivo* ^63^. Similar findings have been observed in the context of CAG repeats in *in vitro* experiments showing that transcription markedly increases the instability of long CAG repeats ^41^. In HD mouse models, correlative *in vivo* evidence also supports our finding that SI and transcription are related. Specifically, instability is correlated with markers of transcriptional activation across several R6 HD transgenic mouse lines expressing the exon-1 fragment of *Htt* with variable expression levels ^64^, and, consistently, similar instability is not observed in the R6/0 line in which the transgene is present in the DNA but not expressed ^65^. To date, SI has been experimentally manipulated in HD mouse models using a wide range of genetic perturbations ^60,66^, as well as postnatal silencing of MMR genes such as *MSH3* and *FAN1* ^67^. Beyond MMR manipulations, direct binding of slipped strand CAG structures by a small molecule (naphthyridine-azaquino-lone) has been purported to stabilize, or even contract, CAG repeats in *Htt* in cells and mouse models ^68^. No reductions in HTT levels were reported with NA treatment, so we believe that we are the first to provide evidence for an intervention that reduces mHTT levels and reduces instability.

We were initially surprised to find that ASO treatment reduces SI in *Htt*’s CAG repeat (Fig. 1). However, multiple mechanisms by which ASOs influence transcription have recently been described ^38–40^. As RNase H is found in both the nucleus and cytoplasm, ASOs can target intronic sequences, and indeed we have developed allele-selective ASOs by largely targeting intronic variants ^31^. One such mechanism occurs after ASO binding to target pre-mRNA, yielding pre-mRNA cleavage and subsequent digestion by nucleases that degrade uncapped mRNA in the nucleus ^38,39^. Exonuclease progression along the pre-mRNA can “torpedo” RNA polymerase off nascent transcripts, thereby reducing transcription downstream of the ASO target. In a more recently described mechanism, chronic treatment with a splice modulating ASO designed for SMA induces repressive chromatin at ASO target sites, suggesting there are at least two distinct mechanisms by which ASOs selectively modulate transcription at targeted transcripts ^40^.

This study was conducted to establish the relationship between HTT-lowering treatments and SI *in vivo*. We find that nuclear-acting HTT-lowering methods, including ASOs, the *lac* repressor, CRISPR-mediated deletion of *HTT*’s promoter, and repressive ZFPs, can reduce SI, while cytoplasmic-acting treatments (e.g. siRNA) do not. Finally, we establish that non-repressive forms of ZFPs that contain only the DNA binding domain reduce SI despite not reducing levels of mHTT, offering a novel approach to treatments in multiple repeat expansion disorders in which SI plays a role.

## Materials and Methods

Detailed methods and protocols are available, along with raw data and code used for statistics and to generate figures, in an online repository (Dryad.org) at the permanent digital object identifier DOI:2010.5061/dryad.0k6djhb98.

### Mice

All mice in the ASO treatment cohorts were group housed in the Western Washington University (WWU) vivarium with *ad libitum* access to food and water. *Htt^Q^*^111^*^/+^* mice (JAX:003598) in the 5.5-month treatment cohort were bred at WWU and *Htt^Q^*^111^*^/+^* mice in the 8-month and 12-month treatment cohorts were acquired from Jackson Laboratories (Bar Harbor, ME) at 1.5 months of age. Beginning at ∼60 days of age, female *Htt^Q^*^111^*^/+^*mice were intraperitoneally (IP) injected with 50 mg per kg body weight (mpk) per week of pan *Htt* targeted ASO (‘*Htt* ASO’), a control ASO without a target in the mouse genome (‘Control ASO’), or 4 µL/g body weight of saline alone. ASOs were synthesized by Ionis Pharmaceuticals (Carlsbad, CA). Sequence information and chemistry of both the *Htt* and control ASOs were published previously ^16^. All procedures performed at WWU were approved by the institutional animal care and use committee (protocol 16-011). *Htt^LacO-Q^*^140^*^/+^* ^34^; *b-actin-LacI^R^ tg* mice (hereafter *Htt^LacO-Q^*^140^*^/+^* mice) were housed at Psychogenics with *ad libitum* access to food and water. For maintaining normal levels of mutant HTT expression, IPTG (2.4 mg/mL) was added to the drinking water of dams pregnant with *Htt^LacO-Q^*^140^*^/+^* pups starting at embryonic day 5. At weaning *Htt^LacO-Q^*^140^*^/+^*mice were treated continuously with IPTG (2.4 mg/mL) in drinking water for the duration of the experiment (6 or 12 months of age) or until either 2 or 8 months of age. Approximately equal numbers of male and female mice were used to quantify somatic expansion. All procedures performed at Psychogenics were approved by the institutional animal care and use committee (protocol 271-0315). All mice in the GalNAc-siRNA treated cohorts were housed and maintained at University of Massachusetts Chan Medical School under IACUC protocol 202000010. For the GalNAc-siRNA experiments, *Htt*^Q111/+^ mice were subcutaneously injected with 10 mg/kg of non-targeting control (NTC) or Htt GalNAc-conjugated siRNA monthly for 4 months beginning at 2 months of age with takedowns at 6 months of age. siRNA were synthesized at UMass Chan Medical School (Worcester, MA) using previously validated sequences (PMID: 31375812) (see full methods below). At takedowns, livers were collected and flash-frozen for downstream processing. For ZFP experiments, *Htt^Q^*^111^*^/+^*mice were housed at Evotec Hamburg in Eurostandard Type II long cages and given access to food and water *ad libitum*. Animal handling was carried out in accordance with the regulations of the German animal welfare act and the EU legislation (EU directive 2010/63/EU). The study protocol was approved by the local Ethics committee of the Authority for Health and Consumer Protection of the city and state of Hamburg (*“Behörde für Justiz und Verbraucherschutz, BJV,* Hamburg).

Crosses between Htt^Q111^ and Htt^ex4/5^ knockout mice were performed at the Massachusetts General Hospital (MGH) in accordance with the recommendations in the Guide for the Care and Use of Laboratory Animals, NRC (2010). All animal procedures were carried out to minimize pain and discomfort, under approved IACUC protocols of the MGH. Animal husbandry was performed under controlled temperature, humidity and light/dark cycles. *Htt*^Q111^ mice on a CD1 genetic background ^10^ were crossed Httex4/5 on a CD1 background ^54^ and male and female heterozygous *Htt^Q^*^111^*^/+^* and hemizygous *Htt^Q^*^111^*^/ex^*^4^*^/^*^5^ (*Htt^Q^*^111^*^/-^*) mice were aged to 3, 6 or 10 months of age.

### GalNAc-conjugated siRNAs

Oligonucleotides were synthesized on a MerMade 12 synthesizer using phosphoramidite solid-phase synthesis. 2′-fluoro RNA or 2′-O-methyl RNA phosphoramidites with standard protecting groups, as well as the 5′-(*E*)-vinyl tetraphosphonate (pivaloyloxymethyl), 2′-*O*-methyl-uridine 3′-CE phosphoramidite (VP) purchased from ChemGenes (Wilmington, MA) or Hongene Biotech (Union City, CA). The antisense strand was synthesized on a UnyLinker 500 Å. The phosphoramidites were prepared at 0.1 M in anhydrous acetonitrile (ACN), except for the 2′-O-methyluridine phosphoramidite, which was dissolved in anhydrous ACN containing 15% anhydrous N,N-dimethylformamide (DMF). Detritylation was performed with 3% trichloroacetic acid in dichloromethane (AIC). Phosphoramidites were activated using 5-(benzylthio)-1*H*-tetrazole (BTT) at 0.25 M in anhydrous ACN, and the coupling time was 4 min. CAP A (20% 1-methyl-*1H*-imidazole in ACN) and CAP B (30% 2,6-lutidine and 20% acetic anhydride in ACN) were used to cap unreacted sites. Oxidation was performed using a solution of 0.05 M iodine in pyridine-water (9:1, v/v; Apex Industrial Chemicals (AIC), Aberdeen, UK) for 4 min. Post-synthesis columns were washed with 10% N,N-diethylethanamine (DEA) in anhydrous ACN.The antisense strand was deprotected using a 10% DEA in ammonium hydroxide solution at 35 °C for 20 hours. The GalNAc sense strands were deprotected using a solution of ammonium hydroxide and methylamine (1:1) for 2 hours at 25 °C. Oligonucleotides were evaporated to dryness before being purified by ion-exchange HPLC (Agilent 1200 PREP system). Oligonucleotide masses were confirmed using LC-MS (Agilent 6530, QTOF). Quantification of single strands was performed on a Nanodrop system. The siRNAs were annealed by adding equal parts of antisense and sense strands, then heating to 95°C and allowing to cool slowly to room temperature.

### ZFPs

#### AAV vector construction and production

ZFP30645, targeting the CAG repeat domain, was obtained from Sangamo (termed ZFP-D in original publication ^30^) and subcloned via EcoRI/HindIII into in the AAV vector pAAV-6P-SWB under the control of the human synapsin1 promoter (p_hSyn1_). A C-terminal HA tag was included to allow for IHC evaluation of the expression levels. Control versions expressing only the KRAB domain or DNA-binding domain were cloned using the same method. Recombinant AAV2/1+2 particles were produced in HEK293 cells co-transfected with AAV vector carrying the transgene and plasmids containing rep and cap genes (pDP1rs and pDP2rs, Plasmid Factory). Producer cells were lysed 48 hours post-transfection, AAV particles released by three freeze-thaw cycles. AAV was purified by iodixanol density centrifugation and a heparin affinity column. Final AAV particles were dialyzed against AAV Storage buffer (10 mM phosphate buffer + 180 mM NaCl + 0.001% Pluronic-F68), aliquoted and stored at -80°C. AAV titers were quantified by qPCR and purity was analyzed by Sypro Ruby gel and endotoxin levels measured by EndoZyme® II Recombinant Factor C Endotoxin Detection Assay. Prior to *in vivo* application, ZFP-expressing AAV vectors were tested *in vitro* in primary cortico-striatal neurons from *Htt^Q^*^111^*^/+^*mice for functionality, i.e. selective mutant Htt downregulation and functional titers confirmed.

#### AAV-ZFP in vivo striatal injections in mice

*Htt^Q^*^111^*^/+^*mice at 2 months of age received bilateral intra-striatal injections. Group 1 received DBD only in the left hemisphere and full ZFP (KRAB+DBD) in the right hemisphere; group 2 received DBD-only in the left hemisphere and KRAB-only in the right hemisphere. Mice were treated with analgesic and individually anesthetized with isoflurane and placed in a stereotaxic instrument (Kopf, Model No. 942). Anesthesia was maintained throughout the surgical procedure. A small hole corresponding to the striatal injection site was made in the skull using an electrical drill (Foredom; Model No. H.30). The coordinates measured according to the mouse bregma were 1 mm anterior, 2.31 mm lateral on right, and 3.6 mm deep (with an angle of 5°) from bregma with flat skull nosebar setting. A total volume of 4 µL per hemisphere (in total 8E9 GCs) rAAV vectors were administered using a Hamilton gas tight syringe (model 1801 RN; Cat. No. 7659-0, customized gauge 26 needle) connected to an automated microinjection pump at a constant flow rate of 500 nl/min. After injection, the surgical wound was sealed, and the animals were kept in a pre-heated home cage until fully recovered.

#### Tissue sampling

Animals were equally assigned to either fresh-frozen or PFA-perfused groups for post-mortem assessment. For the fresh-frozen group, brains were removed and frozen in isopentane at -35°C and subsequently preserved at -80°C. For the PFA-perfused group Mice were deeply anesthetized by intraperitoneal injection of ketamine/xylazine mixture (120 mg/15 mg per kg in 15 μl/g body weight) and transcardially perfused with 80 ml of ice-cold PBS followed by 80 ml of 4% paraformaldehyde using a peristaltic pump. Brain samples were removed, post-fixed overnight at 4°C, and cryoprotected in 30% sucrose solution until saturated. Whole brains were embedded in TissueTek and stored at -80°C.

### CAG Sizing PCR and Fragment Analyses

Genomic DNA from *Htt^Q^*^111^*^/+^* mice was extracted from 10-30 mg of tissues via phenol-chloroform extraction. Genomic DNA concentrations were determined using the Synergy 2 Plate Reader (BioTek). CAG repeat-spanning *HTT* PCR products from both the *Htt^Q^*^111^*^/+^* and *Htt^LacO-Q^*^140^ cohorts were amplified from genomic DNA using 6-FAM-labeled CAG1 forward primer (5’ ATGAAGGCCTTCGAGTCCCTCAAGTCCTTC 3’) and Hu3 reverse primer (5’ GGCGGCTGAGGAAGCTGAGGA 3’) ^69^ using 200 ng genomic DNA. *Htt* amplicons were concentrated from 80 µL to 20 µL using the GeneJET Purification Kit (Thermo Fisher Scientific, cat no. K0702) and eluted in 20 µL of water. Concentrated amplicons were sent to Genewiz (Azenta Life Sciences) for fragment analysis.

### Somatic Instability Analyses

PCR product size distributions from all cohorts were analyzed using the open-source software Peak Scanner 2 or GeneMapper v5 (both produced by Thermo Fisher Scientific). Expansion indices were calculated following previously described methods ^36^, but in our analyses we defined the main allele as the modal allele on a per sample basis, rather than the peak with the greatest fluorescence in tail PCR products from matched mice. Under this definition, CAG distances were determined based on the distance from the modal allele within each sample and expanded alleles were considered peaks with positive CAG distances. To measure modal CAG repeat length and instability index ^36^, data was exported and analyzed with a custom R script, using an inclusion threshold of 10% of modal peak height, and confirmed manually. Linear regression models were built in R and compared by an analysis of variance (ANOVA).

### Western blotting

Protein was extracted from 40-55 mg of tissue in RIPA buffer (150 mM NaCl, 1.0% NP-40, 1% sodium deoxycholate, 0.1% SDS, 25 mM Tris-HCl) containing protease and phosphatase inhibitors (Thermo Fisher, 78441). Frozen tissues were mechanically homogenized using the Bead Blaster 24 (Benchmark) at room temperature for 2 cycles at 7m/s with cycles consisting of 2 x 30 seconds with a 10 second intermission. Following homogenization, samples were centrifuged for 15 minutes at 15k*g to clear and protein concentration was determined by Pierce BCA assay (Thermo Fisher Scientific, cat no. 23225) according to the manufacturer’s protocol. Denatured protein lysates were loaded into 3-8% Tris-Acetate mini gels (Thermo Fisher Scientific), separated electrophoretically, and transferred to a PVDF membrane (Millipore) for 17-20 hours at 4°C. Membranes were incubated overnight at 4°C in primary antibodies targeting HTT. Antibodies used were monoclonal, rabbit anti-huntingtin antibody EPR5526 (Abcam ab109115) 1:1,000 in Intercept blocking buffer (Li-cor, 927-60001) plus 0.05% tween and 1:15,000 goat anti-rabbit secondary antibody (Li-Cor, 926-32211) diluted in Intercept blocking buffer, 0.05% tween, and 0.01% SDS. Membranes were imaged using the Li-Cor Odyssey Fc Instrument. Signal intensity was quantified in the ImageStudio Lite software, normalized to the total protein signal of the corresponding lane, and expressed relative to the average signal of saline-treated mice within the same blot.

### qRT-PCR

Tissue (30-50 mg) was homogenized in 1 mL of Qiazol lysis reagent using the Bead Blaster 24 (Benchmark). RNA was extracted using the RNeasy Lipid Tissue Mini kit (Qiagen 74804) according to the manufacturer’s protocol. cDNA synthesis and qRT-PCR was conducted using a primer and probe set for *Htt* (Thermo Fisher, Mm01213820_m1) that detects both the *Htt^Q^*^7^ and *Htt^Q^*^111^ alleles. RNA concentration and quality was quantified on the NanoDrop Spectrometer and 0.5 to 2 μg of RNA was synthesized to cDNA using the SuperScript III First-Strand Synthesis System (Thermo Fisher Scientific), per manufacturer’s instructions. qRT-PCR was carried out with TaqMan Universal Master Mix (Thermo Fisher Scientific, 4440042) with the following reaction components; 7.5 μL master mix, 3 μL dH_2_O, 3 μL cDNA, and 1.5 μL probes. The reaction was quantified on the QuantStudio 5 ProFlex system. Data were analyzed using the comparative Ct (cycle threshold) method (ΔΔCt). The reference genes *ActB* (Thermo Fisher, Mm02619580_g1) and *Tfrc* (Thermo Fisher, Mm00441941_m1) was used to determine the relative expression of *Htt* (Hs00918174_m1). This was expressed relative to the average expression in the control samples. Fold change was given by 2^ΔΔCt^. WT and mutant *Htt* knockdown in striata of ZFP injected mice was assessed by qPCR using the following primers: forward 5’ CAGGTCCGGCAGAGGAACC 3’ and reverse 5’-TTCACACGGTCTTTCTTGGTGG-3’ for WT, and forward 5’-GCCCGGCTGTGGCTGA-3’ and reverse 5’-TTCACACGGTCTTTCTTGGTGG-3’ for mutant *Htt*, respectively. cDNA was generated using SuperScript™ III First-Strand Synthesis System (Thermo Fisher Scientific), quantitative PCR was performed using SYBR Green I Master Mix (Roche) and LightCycler 480 II real-time PCR instrument (Roche). qPCR primers for gene expression analysis are from Metabion. 10 µl qPCR reactions were used for qPCR: 5 µl SYBR Green I Master Mix (Roche), 0.5 µl of 10 µM forward and reverse primer (final concentration 0.5 µM each primer), 3 µl H_2_O and 1 µl of cDNA sample. Thermal cycling conditions were as follows: 1) 95°C for 10 min, 2) 95°C for 5 sec, 3) 56°C for 5 sec, 4) 72°C for 5 sec, 5) Plate Read, 6) Repeat steps 2 to 5, 45 times, and 7) Melting curve analysis 65°C to 95°C.

### Pre-mRNA qPCR

RNA was extracted and cDNA synthesized as described above. For nascent RNA extractions, the Click-iT kit (Invitrogen C10337) was used according to the manufacturer’s protocol in homozygous STHdhQ111 cells (RRID:CVCL_M591). Pre-mRNA was measured using three intron-exon spanning primer sets along the *Htt* transcript: exon 2-intron 2 (Forward 5’-ACTCTCAGCCACCAAGAAAGAC-3’, reverse 5’-AGATGGCTCAGCACTGGCTAC-3’), intron 37-exon 38 (forward 5’-TGCGAGGATGATGGTGATTGT-3’, reverse 5’-TCCACAGGACGTAGGGAGG-3’), and exon 66-intron 66 (Forward 5’-GCCTCTCCTGCTTCCTTGTTA-3’, reverse 5’-ACCTGAACTGGATGCTAAGTGAA-3’). The reference gene *Tfrc* (forward 5’-TAGGCCACCTTAGTGACTGC-3’, reverse 5’-GTCTCCACGAGCGGAATACA-3’) was used to determine the relative expression of *Htt* pre-mRNA. Data were analyzed using the modified comparative Ct (cycle threshold) method (ΔΔCt) and fold change was given by (primer efficiency*0.02)^ΔΔCt^ ^70,7172^. This was expressed relative to the average expression in the control samples.

### bDNA

#### DNA and RNA co-isolation from fresh-frozen striatal punches of ZFP transduced Htt^Q111/+^ mice

Four punches from frozen striata were harvested from coronal frozen sections of 0.5 mm height: two punches with a diameter of 1.5 mm and 0.5 mm height close to the injection site and two punches with a diameter of 1 mm and 0.5 mm height adjacent to the first punches and used for DNA-RNA co-isolation. DNA and RNA co-isolation of mouse striatal punches were prepared with the AllPrep DNA/RNA 96 kit AllPrep DNA/RNA Kit (Qiagen, #80284) according to the manufacturer’s instructions. Samples were first disrupted in lysis buffer via bead beating using Precellys CK14 Lysing kit (Bertin corp, Ref. no. P000933-LYSK0-A). An additional on-column DNase digest for RNA samples was performed after the RNA binding step and before the first wash step. On-column DNase digestion was performed with RNase-free DNase Set (Qiagen Cat. No. 79254) according to the manufacturer’s instructions with a 15 minute incubation at RT.

#### bDNA assay

QuantiGene Plex was developed by ThermoFisher and is a hybridization-based assay using the branched DNA (bDNA) technology, which uses signal amplification rather than target amplification for direct measurement of RNA transcripts. Processing and analysis of samples were performed following the manufacturer’s instructions using purified RNA.

To assess the linearity of the 15-plex QuantiGene probe set, 400 ng of purified RNA was used as the starting material for a 2-fold serial dilution. Linearity was confirmed across all targets and the maximum input was selected for analysis of experimental samples. Each sample was run in duplicate. Specifically, 20 µL of purified RNA was mixed with 80 µL of Working Bead Mix containing the 15-plex probe set, capture beads, and blocking reagents. This mixture was incubated overnight at 54°C in a hybridization plate while shaking in a VorTemp incubator. After incubation, pre-amplifier, amplifier, and biotinylated probe reagents were sequentially added while shaking at 50°C to ensure signal amplification.

Finally, streptavidin R–phycoerythrin conjugate (SAPE) was added, generating a fluorescent signal associated with individual beads and, consequently, individual genes. The signal was reported as median fluorescence intensity (MFI) and is proportional to the number of mRNA transcripts captured on the beads. Our panel included four housekeeping genes, and the geometric mean of their background-subtracted MFIs was used to normalize the data for each gene.

### MSD

Total HTT levels were quantified using an electro-chemiluminescent ELISA measured with a MESO QuickPlex SQ120MM (Meso Scale Discovery; MSD) according to previously described methods ^70,71^. Tissues were homogenized in a non-denaturing lysis buffer (150mM NaCl, 20mM Tris-HCl, 1mM EDTA, 1mM EDTA, 1% v/v Triton X-100, 0.01M NaF, 1mM PMSF, 1× protease and phosphatase inhibitors (Sigma P5726, Sigma P0044, Roche 11836170001) in tubes containing 1.4mm zirconium oxide beads at 6 m/s in three 30 second intervals with 5 minutes on ice between rounds. Lysates were centrifuged for 20 minutes at 20,000 x g at 4C. Supernatant was transferred to a new tube and centrifuged again for 20 minutes at 20,000 x g at 4C, then transferred to a second new tube. 96-well MSD plates (MSD, L15XA-6) were coated with capture antibody (CHDI-90002133, 8ug/mL) in carbonate-bicarbonate coating buffer for 1 hour with shaking at 750 RPM. Plates were then washed 3 times with wash solution (0.2% Tween-20 in PBS) and blocked with blocking buffer (2% BSA, 0.2% Tween-20 in PBS) for 1 hour with shaking, then washed 3 more times. Samples were diluted (brain: 2ug/uL, liver: 4ug/uL) in 20% MSD lysis buffer, 80% blocking buffer and incubated for 1 hour with shaking. Plates were washed 3 more times, followed by incubation with sulfo-tag conjugated secondary antibody (D7F7-Sulfo-Tag, 1:1800) for 1 hour with shaking. Plates were washed a final 3 times, and Read Buffer B (MSD, R60AM-4) was added to the plate before reading on a QuickPlex SQ 120MM.

### Immunohistochemistry

Coronal sections of 25 µm from ZFP-treated mice were cut using a cryostat and collected as free-floating in 24-well plates. All stainings were performed with free-floating sections. Sections were permeabilized in 0.3% Triton X-100/PBS, blocked in 10% normal goat serum/PBS, and incubated with the primary antibodies diluted in 1% normal goat serum, 0.1% Triton X-100 in PBS at 4°C overnight. Primary antibodies used were anti-NeuN (1:1000, Millipore, ABN90P), polyclonal rabbit anti-HA (C29F4, 1:400, Cell signaling, 3724S), anti aggregated HTT (S830, kindly provided by Gill Bates, UCL), chicken anti-GFAP (1:1000, Abcam, ab4674) or rabbit anti-Iba1 (1:1000, WAKO, 019-19741). After three washes in PBS, sections were incubated with secondary antibodies (anti-Rabbit IgG CF™ 568, Anti-Guinea Pig CF™ 647, and anti-Chicken CF™ 488: all 1:1000, Sigma Aldrich) for 2 hours at room temperature. Subsequently, sections were washed twice in 0.1% Triton X-100/PBS, incubated with DAPI (1:10,000, Sigma Aldrich, D9542), washed once in 0.1% Triton X-100/ PBS, and mounted using 10 mM Tris-buffered saline pH 7.4 in 24-well glass-bottom plates (24 Well SensoPlate^TM^, Greiner, #662892) suitable for imaging with the Opera High Content Screening System (PerkinElmer Inc.). For fluorescence preservation, sections were covered with an aqueous mounting medium (Anti-Fade Fluoroshield Mounting Medium, Abcam, 104135).

Automated image acquisition was conducted using the Opera® High Content Screening system and Opera software 2.0.1 (PerkinElmer Inc.), using a 40× water immersion objective (Olympus, NA 1.15, lateral resolution: 0.32 µm/pixel). Image analysis scripts for HA+/NeuN+ cells were developed using Acapella® Studio 5.1 (PerkinElmer Inc.) and the integrated Acapella® batch analysis as part of the Columbus® system.

## Supplemental Figures and Tables

**Supplemental Figure 1:**
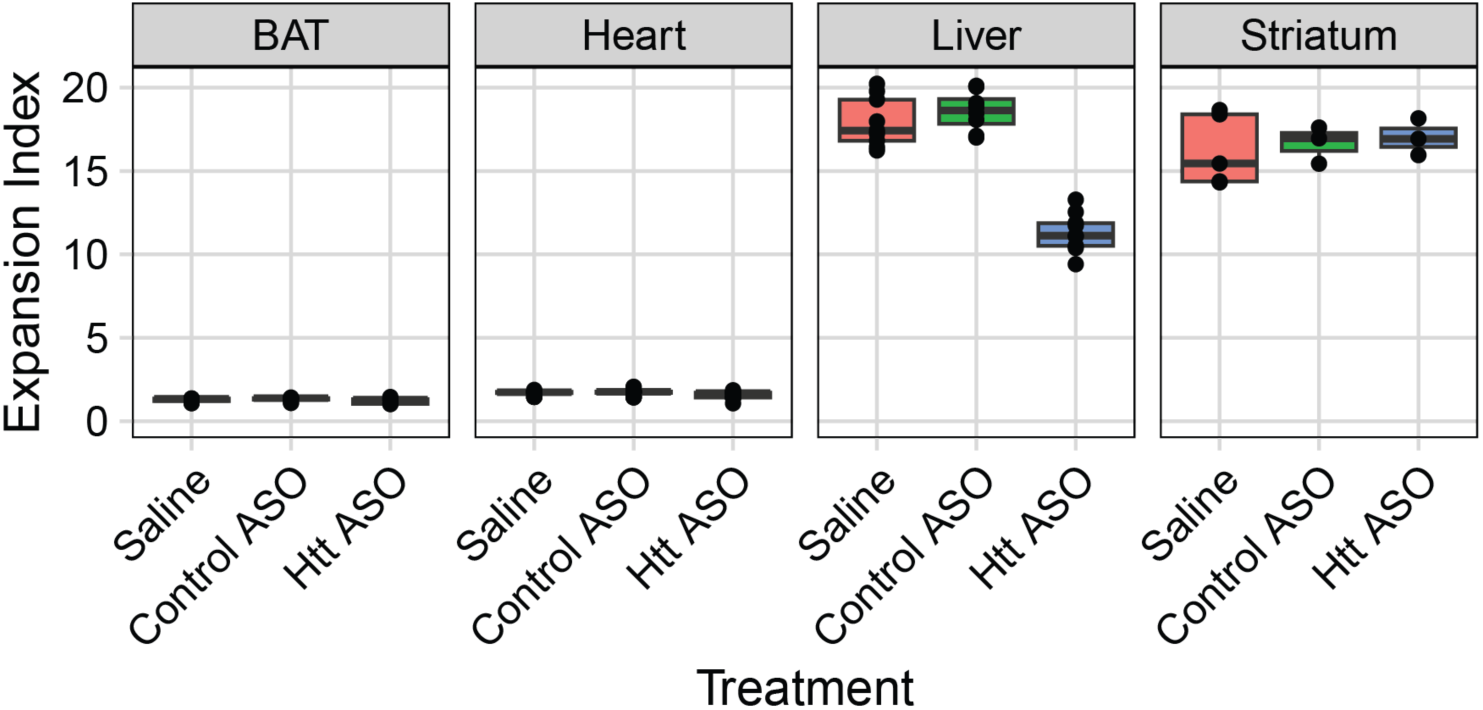
Peripheral ASO treatment does not reduce instability in tissues other than liver. Brown adipose tissue (BAT) and heart are stable tissues with very low baseline instability and are not affected by Htt ASO treatment. Striatum is an unstable tissue, but Htt ASO is dosed peripherally and cannot cross the blood-brain barrier, so instability is unaffected. Liver is both unstable and accessible by the ASO, and its instability is markedly reduced by ASO treatment.

**Supplemental Figure 2:**
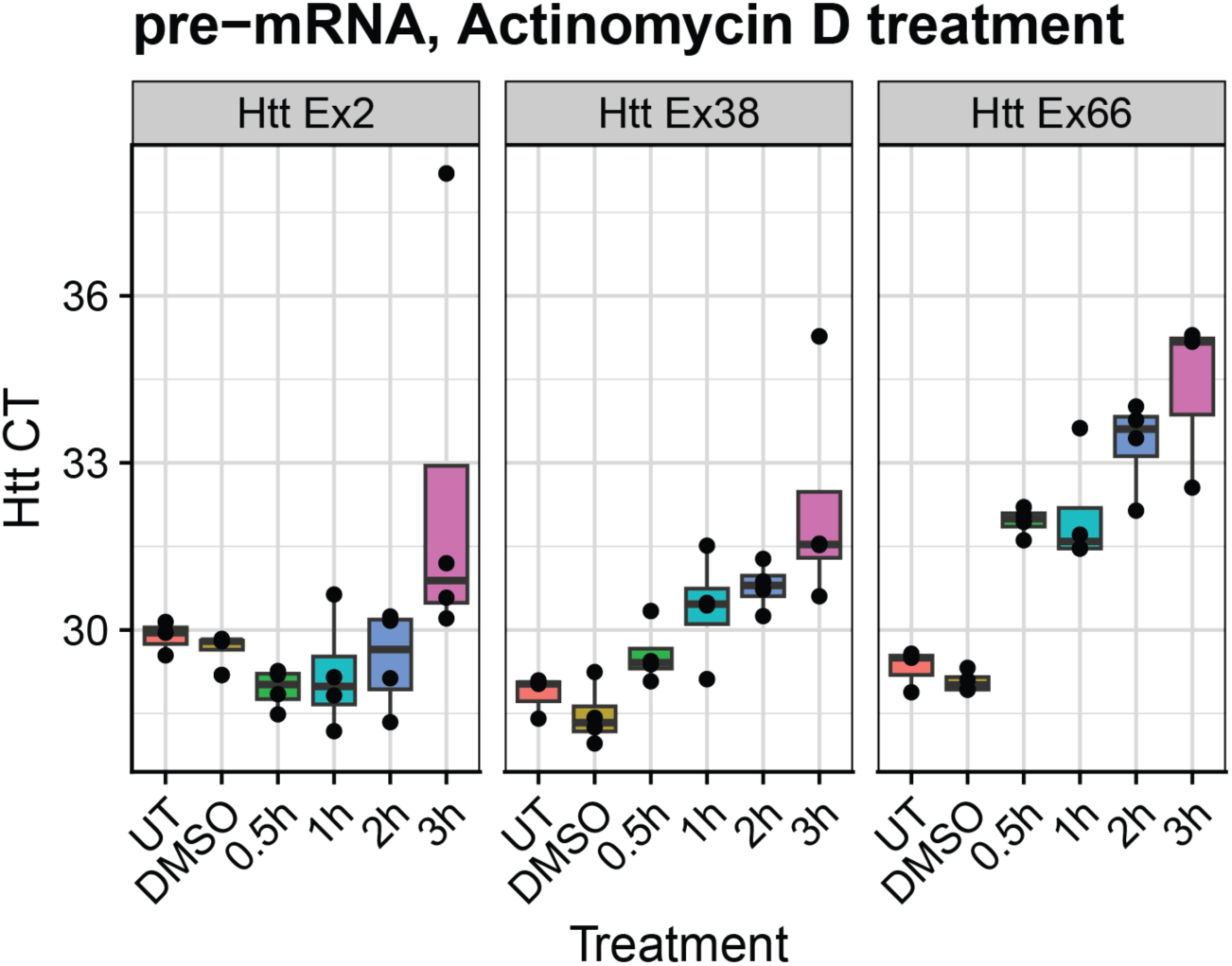
Actinomycin D treatment leads to robust reductions in *Htt* pre-mRNA. In AML12 cells, pre-mRNA qPCR signal for Htt RNA decreases with increasing lengths of treatment with actinomycin D, a known transcriptional inhibitor.

**Supplemental Figure 3:**
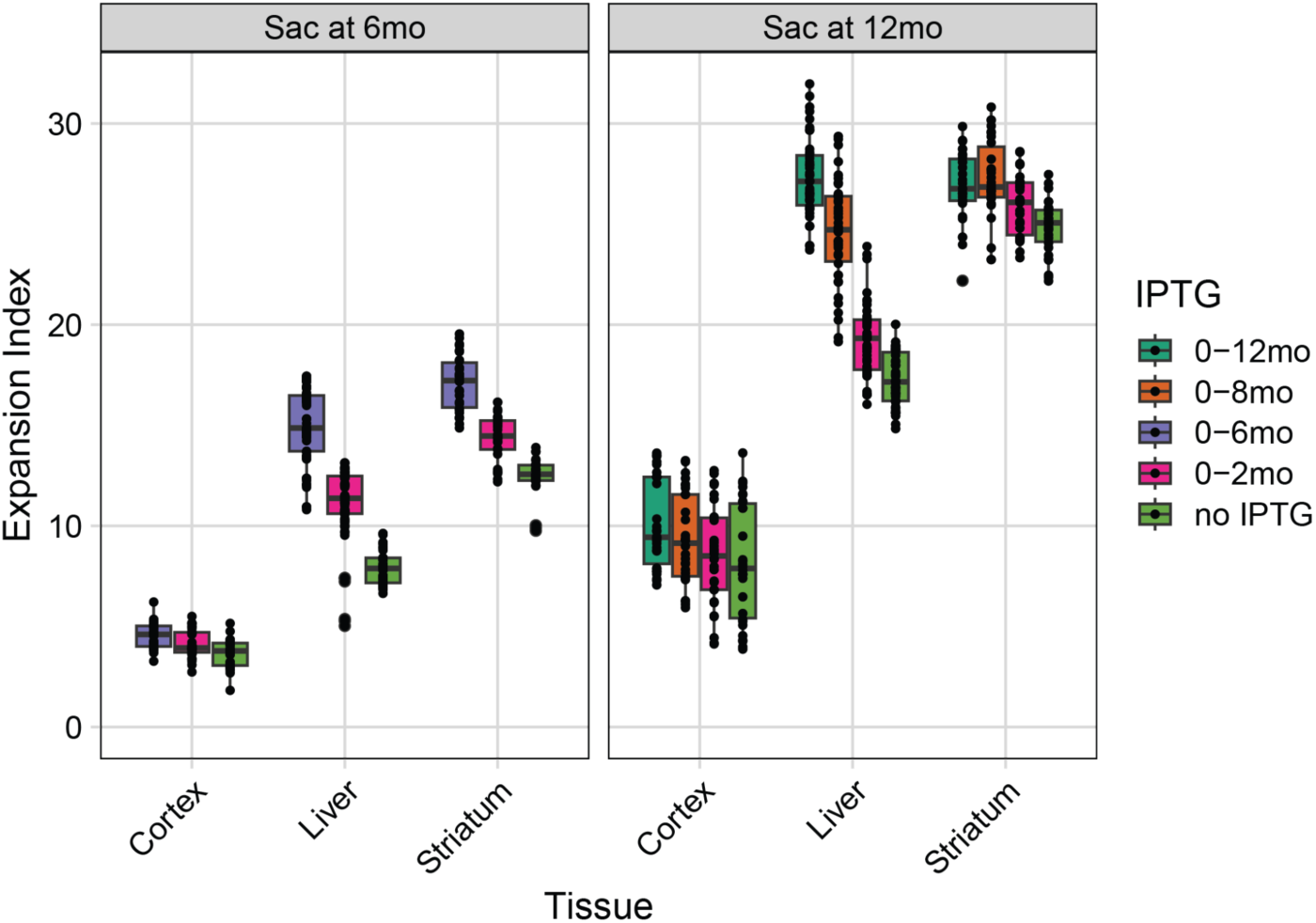
Complete, cross-tissue, LacO dataset. Expansion index decreases with longer windows of *Htt* repression, ie, shorter windows of IPTG treatment. This effect is seen in cortex, liver, and striatum, and in mice sacrificed at both 6 and 12 months of age.

**Supplemental Figure 4:**
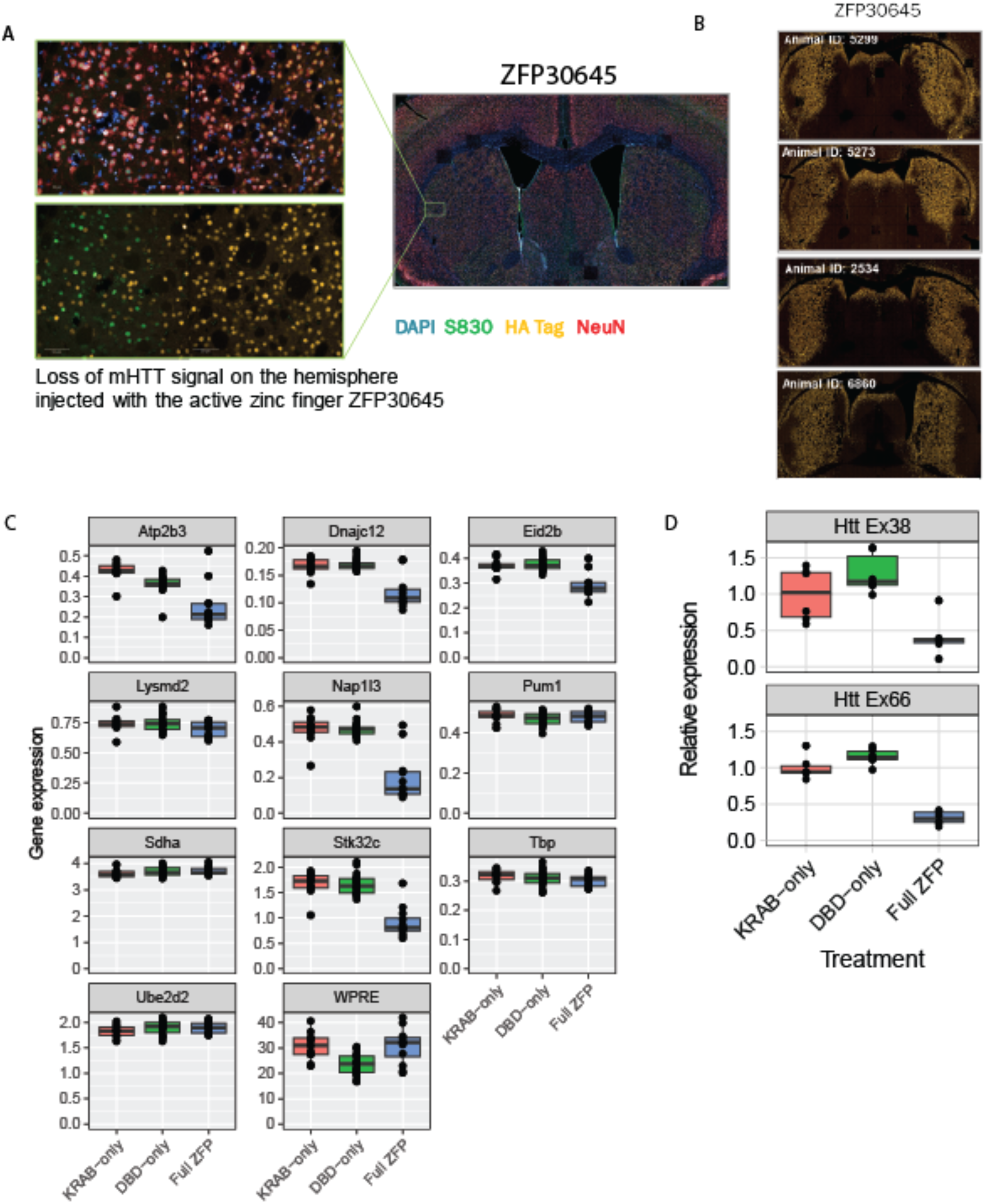
Experimental validation of the delta-DBD and delta-KRAB derivative ZFP constructs. **A)** Immunohistochemical staining showing loss of mHTT in the transduced area. **B)** HA-tag staining showing successful distribution of ZFP into the striatum. **C)** Off-target gene data. **D)** Results of *in vitro* study showing no reduction of transcription in DBD-only construct.

**Supplemental Table 1: Raw data underlying all graphs.**

**Supplemental Table 2: Comparison of inherited CAG sizes across cohorts.**

## Acknowledgments

We would like to thank Tammy Gillis of the MGH Mission Driven Service Core for DNA Fragment Analysis.

## Funding

Funding for this project was provided to JB Carroll via a research agreement between CHDI Foundation and UW and WWU (Record number # A-18222), and by a research agreement between CHDI and UMass Chan Medical School (A-5038). Additional funding was provided by R01 NS049206 and a research agreement between Mass General and CHDI.

## Declaration of Interests

A.N., A.S., J.F., T.S., A.C., I.L. are Employees of Evotec, and may have stock options. C.F.B. and H.K. Are full time employees at, and hold shares in, Ionis Pharmaceuticals. V.C.W. was a founding scientific advisory board member with a financial interest in Triplet Therapeutics Inc., her financial interests were reviewed and are managed by Massachusetts General Hospital (MGH) and Mass General Brigham (MGB) in accordance with their conflict of interest policies. V.C.W. is a scientific advisory board member of LoQus23 Therapeutics Ltd. and has provided paid consulting services to Acadia Pharmaceuticals Inc., Alnylam Inc., Biogen Inc., Passage Bio and Rgenta Therapeutics and has received research support from Pfizer Inc. J.B.C. Has provided paid consulting and/or conducted sponsored research for Wave Life Sciences, Skyhawk Therapeutics, Cajal Neuroscience, Ionis Pharmaceuticals, and Alnylam, and Gudiepoint. D.H., D.M. and T.V.are full-time employees of CHDI Foundation. A.K. discloses ownership of stocks in RXi Pharmaceuticals and Advirna, and is a founder of Atalanta Therapeutics and Comanche Biopharma. R.B. received consulting fees from Takeda.

## Notes

https://doi.org/10.5061/dryad.0k6djhb98

